# Exogenous corticosterone administration during pregnancy alters placental and fetal thyroid hormone availability in females

**DOI:** 10.1101/2023.07.05.547278

**Authors:** Emmanuel N. Paul, Salome Shubitidze, Rodaba Rahim, Imani Rucker, Liana Valin, Stefanos Apostle, J. Andrew Pospisilik, Karen E. Racicot, Arianna L. Smithb

**Author notes:** Correspondence: Arianna L. Smith, Ph.D., Kenyon College, Biology Dept., 202 N College Road, Gambier, OH 43022. Co-Senior Authors (Karen E. Racicot, Arianna L. Smithb). Declarations of interest: none. Whitehead Institute for Biomedical Science, Cambridge, MA 02135.

## Abstract

**Introduction:** Maternal prenatal stress is associated with adverse pregnancy outcomes and predisposition to long-term adverse health outcomes in children. While the molecular mechanisms that govern these associations has not been fully teased apart, stress-induced changes in placental function can drive sex-specific phenotypes in offspring. We sought to identify and examine molecular pathways in the placenta that are altered in response to maternal prenatal stress.

**Methods:** Using a mouse model of maternal prenatal stress, we conducted RNA-seq analysis of whole placenta at E18.5. We used qRT-PCR to validate gene expression changes in the placenta and in a trophoblast cell line. ELISAs were used to measure the abundance of thyroid hormones in maternal and fetal serum and in the placenta.

**Results:** *Dio2* was amongst the top differentially expressed genes in response to elevated maternal stress hormone. *Dio2* expression was more downregulated in female placenta from stressed dams than both female control and male placenta. Consistent with Dio2’s role in production of bioactive thyroid hormone (T3), we found that there was a reduction of T3 in placenta and serum of female embryos from stressed dams at E18.5. Both T3 and T4 were reduced in the fetal compartment of the female placenta from stressed dams at E16.5. Stress hormone induced reduction in thyroid hormone in females was independent of circulating levels of TH in the dams.

**Discussion:** The placental thyroid hormone synthesis pathway may be a target of maternal stress and modulate fetal programming of health and disease of offspring in a sex-specific fashion.

## Introduction

Maternal stress during pregnancy is associated with adverse health outcomes for both the mother and baby. Maternal prenatal stress increases risk for intrauterine growth restriction, preeclampsia and preterm birth [1–3]. It is also associated with long-term adverse health outcomes in offspring, to include asthma, and other immune disorders, and neurocognitive and metabolic disorders [4]. Notably, the incidence of these health outcomes is often dominant in one sex over another [5–11]. Using a mouse model of low-grade prenatal stress during late pregnancy, we previously showed that mice born to dams with elevated stress hormone have heightened responses to airway allergen when compared to controls, and that these responses differ based on sex of the offspring [9].

The placenta is an ephemeral organ essential for the establishment and maintenance of a healthy pregnancy. During its transient existence, the placenta has multiple and critical functions. It serves as a physical and physiological selective barrier that supplies the developing fetus with oxygen and nutriments but limits the exposure to potential harmful environmental pollutants and maternal hormones [12]. In early pregnancy, it serves as a buffer against excess maternally derived stress hormone, cortisol (corticosterone in rodents, CORT) [13]. Near term, transport of CORT across the placenta is required for maturation of fetal organs, particularly the lung [14]. This protective function of the placenta is mediated by the enzyme 11-β-Hydroxysteroid Dehydrogenase (11β-HSD2), which converts active CORT into its inactive form.

Expression of 11β-HSD2 is high early in gestation and is reduced in late pregnancy [15, 16]. Additionally, the placenta expresses the glucocorticoid receptor (GR) which, when bound by cortisol, transcriptionally regulates expression of target genes and mediates the stress response [15, 17, 18]. Thus, the placenta both protects the fetus from and is responsive to maternally derived stress hormone.

The placenta mediates the effects of maternal prenatal stress in a sex-specific manner [5–7]. Expression of GR isoforms differ in the placenta based on both fetal sex and age [17, 19, 20]. Sex-specific differences in the placenta’s global gene expression in response to maternal prenatal stress have been reported in both humans and animal models [21–24]. In a mouse model of maternal prenatal stress in early pregnancy, neurocognitive defects observed specifically in male offspring born to stressed dams were recapitulated by placental knockout of *O-GlcNAc transferase (Ogt)*. Notable, *Ogt* expression was altered only in the male placenta in response to maternal stress [25, 26]. However, timing and duration of maternal stressors appears to be important for placenta-mediated responses and reprogramming effects [27, 28]. Thus, work remains to be done on the cellular and molecular pathways that are altered by maternal prenatal stress and the consequences of these alterations on fetal and offspring health.

Thyroid hormone (TH) signaling during pregnancy has been shown to be sensitive to a variety of maternally experienced environmental insults, including maternal stress [29–32]. Administration of dexamethasone, a synthetic cortisol analog, to pregnant ewes resulted in increased concentrations of the active thyroid hormone, triiodothyronine (T3), in the fetus but reduced T3 and inactive thyroid hormone, thyroxine (T4), in the ewe. Additionally, prenatal administration of dexamethasone to the ewe altered expression of deiodinase (DIO) enzymes in the placenta [29]. These enzymes function to metabolize thyroid hormone via deiodination [33]. In a mouse model of maternal stress where pregnant dams were exposed to predator odor, increased circulating T4 and upregulation of the thyroid hormone receptor α in the brain of adult offspring was observed [31]. Taken together, these results implicate TH biosynthesis and signaling pathways as targets of maternal prenatal stress.

Maintaining the supply of THs (both T3 and T4) during pregnancy is essential for appropriate growth and development of the fetus and disruption is associated with short and long-term adverse outcomes [34]. For example, decreased levels of fetal THs have been reported in cases of intrauterine growth restriction [35]. The placenta has a featuring role in the metabolism and transport of thyroid hormones from the maternal to fetal circulation via expression of DIO enzymes and TH transporters, respectively [36]. Altered expression of genes involved in placenta TH metabolism and transport have been implicated in IUGR, recurrent miscarriage and gestational diabetes [37, 38]. Thus, in addition to its sensitivity to maternal stress, placenta TH signaling is also a regulator of pregnancy health and fetal outcomes.

Here we report the use of a mouse model of maternal prenatal stress to study the effects of elevated stress hormone in late pregnancy on TH bioavailability in the placenta and fetus. We show that *iodothyronine deiodinase II* (*Dio2*), the gene responsible for converting T4 to T3 in tissues, is downregulated in response to maternal CORT treatment during pregnancy and that this downregulation was concomitant with reductions in placental and fetal T3 in females.

## Methods

### Animal treatments

Nulliparous C57BL6/J mice were purchased from Jackson Laboratories (Bar Harbor, ME, USA). Male and female mice were housed separately under controlled room temperatures (22 ± 2 °C) with a 12 h dark/light cycle and *ad libitum* access to food and water. Ethical approval for all animal experimentation was obtained by the Institutional Animal Care and Use Committee (IACUC) at Michigan State University and Kenyon College.

Eight – to - sixteen week old females were placed with males overnight (12h). The following morning, mating was confirmed by the presence of a vaginal plug, and this time-point was considered to be embryonic day 0.5 (E0.5). To model the effects of chronic, mild stress, pregnant dams were administered corticosterone (CORT) in drinking water as previously described [9]. Briefly, beginning at E12.5, pregnant mice were administered vehicle (0.1875% 2-hydroxy-β-cyclodextrin) or 50 µg/mL CORT (Sigma-Aldrich, St. Louis, MO) in distilled drinking water (Figure 1A).

**Figure 1:**
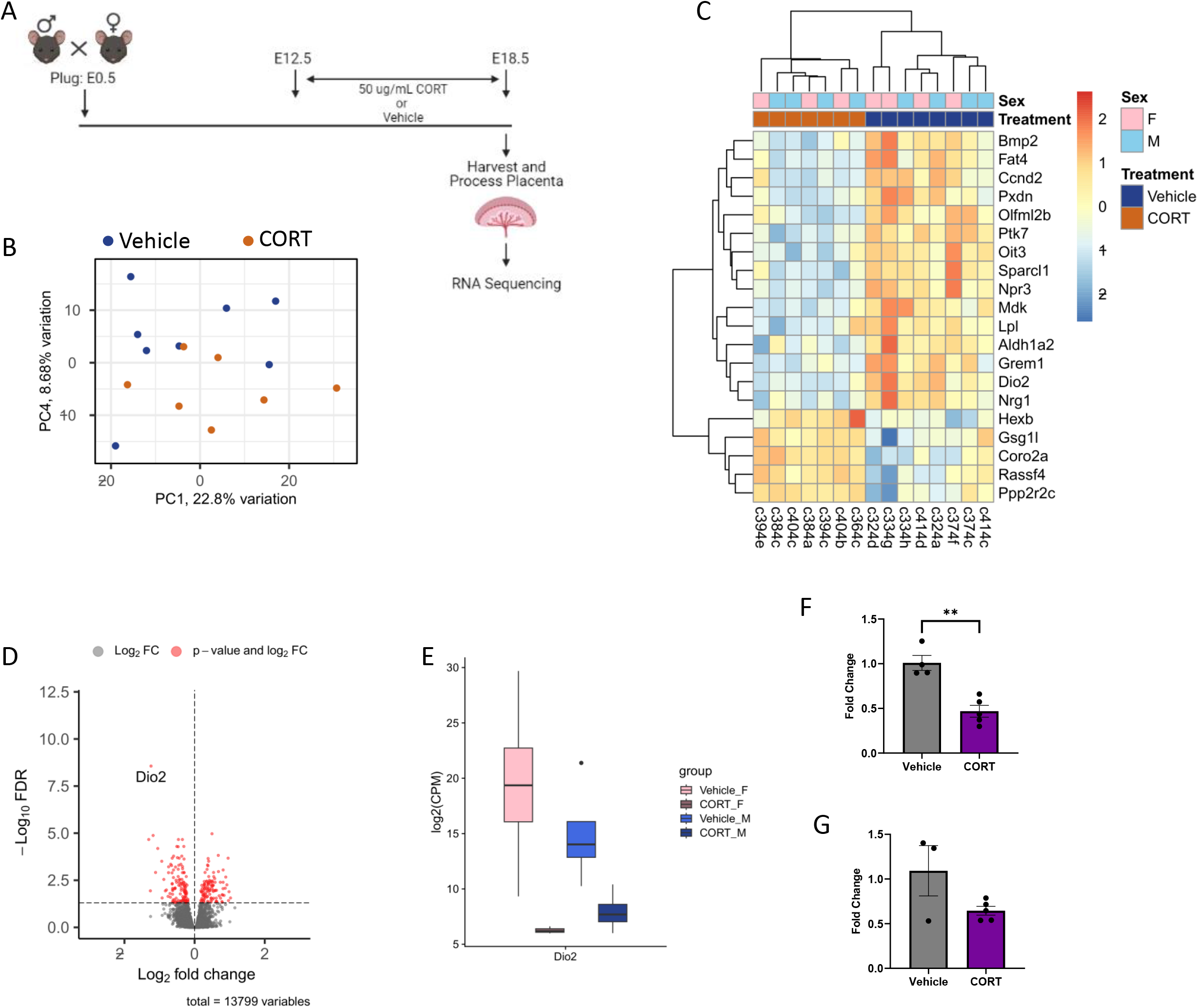
Differential gene expression in placenta from Vehicle and CORT-treated dams at E18.5. A) Mouse mating treatment and collection strategy. B) Principal component analysis of placental transcriptomes. C) Heatmap of top 20 differentially expressed genes with FDR <0.05. n = 8 placenta from 4 dams/treatment – one male and one female placenta were selected from a single dam. D) Volcano plot showing 238 differentially expressed genes (red) at FDR <0.05. Dio2 is indicated. E. Box plot showing *Dio2* expression as log_2_(CPM) in placental samples stratified by sex and treatment. F-G) qPCR validation of *Dio2* expression in female (C) and male (D) whole placenta. N = 4-5 placenta from independent dams. Gene expression was normalized to beta-actin and set relative to the Vehicle group. Data are presented as average fold change (2^-ddCt) +/-SEM. Statistical analysis was performed on dCt values (see methods). **p <0.01, unpaired t-test. Fig 1A created with BioRender.com

### Tissue collection

Pregnant dams were sacrificed at E16.5 or E18.5. Whole blood from pregnant dams and individual fetuses was collected at the time of euthanasia via cardiac puncture and trunk bleed, respectively. Whole blood was spun at 1,000xg for 10 minutes at room temperature and serum was collected and stored at -80℃ until use. Fetal serum was pooled by sex within a litter. Tail snips were collected from each fetus, stored at -80℃ and used to determine fetal sex.

Whole placenta were harvested from euthanized dams at E16.5 or E18.5 and flash frozen in liquid nitrogen. To collect trophoblasts (fetal portion of the placenta) at E16.5, whole placenta were separated from the implantation site and the maternal decidua was removed manually using forceps [39]. The remaining, fetal portion of the placenta was flash frozen in liquid nitrogen. Placental tissues were stored at -80℃ until use.

### Determination of fetal sex

Embryo sex was determined after collection via SRY genotyping (see Supplementary Table 1) [40]. Briefly, DNA was extracted from tail snips using the Zymo Quick-DNA Miniprep Plus kit (Irvine, CA USA). Multiplex PCR was conducted to detect the presence of SRY locus and an internal control sequence using the Phire Tissue Direct PCR Master Mix (Thermo Scientific, Waltham, MA USA). Cycling conditions were as follows: 95℃ for 10 minutes, 34 cycles of 94℃ for one minute, 50℃ for one minute, and 72℃ for one minute, followed by a final extension step of 72℃ for five minutes. PCR products were resolved on a 1.5% agarose gel.

### RNA Isolation, Library Preparation and Sequencing

Total RNA was isolated from 16 placentas (3-4 placentas/fetal sex/group) using an RNAeasy kit (Qiagen, Venlo, The Netherlands). Isolated RNA was stored at −80 °C in nuclease-free water. Nanodrop 1000 spectrophotometer (Thermo Fisher Scientific, Fairlawn, NJ, USA) and Agilent 2100 Bioanalyzer (Agilent Technologies, Santa Clara, CA, USA) instruments were used to measure RNA concentration and quality, according to the manufacturers’ protocols. Libraries were prepared by the Van Andel Genomics Core from 1ug of total RNA using the KAPA stranded mRNA kit (v5.16) (Kapa Biosystems, Wilmington, MA USA). RNA was sheared to 300-400 bp. Prior to PCR amplification, cDNA fragments were ligated to Bioo Scientific NEXTflex dual adapters (Bioo Scientific, Austin, TX, USA). Quality and quantity of the finished libraries were assessed using a combination of Agilent DNA High Sensitivity chip (Agilent Technologies, Inc.), QuantiFluor® dsDNA System (Promega Corp., Madison, WI, USA), and Kapa Illumina Library Quantification qPCR assays (Kapa Biosystems). Individually indexed libraries were pooled and 75 bp, single end sequencing was performed on an Illumina NextSeq 500 sequencer using a 75 bp HO sequencing kit (v2) (Illumina Inc., San Diego, CA, USA), with all libraries run across 2 flowcells. Base calling was done by Illumina NextSeq Control Software (NCS) v2.0 and output of NCS was demultiplexed and converted to FastQ format with Illumina Bcl2fastq v1.9.0.

### RNA-Seq Analysis

Reads from 16 placentas, were trimmed for quality and adapters using TrimGalore (version 0.6.5), and quality trimmed reads were assessed with FastQC (version 0.11.7). Trimmed reads were mapped to the mouse genome (GRCm38.p6) using STAR (version 2.7.9a). Reads overlapping Ensembl annotations were quantified with STAR prior to model-based differential expression analysis using edgeR-robust. Genes with low counts per million (CPM) were removed using the filterByExpr function from edgeR. Scatterplots of two selected principal components was constructed with the pca function of the PCAtools R package (version 2.5.13) to verify sample separation prior to statistical testing. Generalized linear models were used to determine if principal components were significantly associated with cell type. Genes were considered differentially expressed if their respective edgeR-robust false discovery rate (FDR) corrected p-values were less than 0.05. Differential expression was calculated by comparing CORT versus Vehicle placentas. Differentially expressed genes (DEG) were visualized with volcano plots and heatmaps generated using the EnhancedVolcano (version 1.8.0) and pheatmap (version 1.0.12) packages in R. Box plots of the log_2_(CPM) values were generated using the R package ggplot2 (version 3.4.0).

T3 and T4 Analysis:

Total T3 and T4 were detected in serum and placental lysates using the Mouse Triiodothyronine (NBP2-60183-1KIT) and Mouse Thyroxine ELISA Kit (NBP2-60162-1KIT; Novus Biologicals, Littleton, CO). Placental homogenates were produced according to manufacturer’s protocol. Briefly, whole placentae or trophoblasts were homogenized in ice-cold 1X PBS using a probe homogenizer. Homogenization was followed by two freeze-thaw cycles where homogenates were frozen rapidly at -80℃ then thawed at rapidly at 37℃. The homogenate was cleared by centrifugation at 5000 × g for 5 minutes and supernatants were assayed immediately. Placental homogenates and fetal and maternal serum were assayed for both T3 and T4 without dilution according the manufacturer’s protocol. The resulting absorbance values was analyzed using 4-PL to determine the concentration of T3 or T4 in each sample. Total protein was quantified in placenta/trophoblast homogenates using the Pierce BCA Protein Assay (Thermo Fisher Scientific, Waltham, MA). T3 and T4 concentrations was normalized to total protein for each sample.

Cell Culture and Treatments:

The first-trimester extravillous trophoblast cell line, Sw.71 cells (kind gift from Dr. Gil Mor), were cultured in DMEM-F12 1:1 supplemented with 10% FBS and 10IU/mL penicillin/streptomycin antibiotic cocktail at 37℃, 5% CO2. For gene expression analysis, 300,000 cells/well were plated into 6-well plates. Cells were treated with 500 ng/mL hydrocortisone (HC, Sigma-Aldrich, St. Louis, MO), 1uM RU486 for 6- or 24-hours. After incubation, cells were harvested using 0.05% trypsin-EDTA (Life Technologies,Carlsbad, CA) and cell pellets were stored at -80℃ until use.

RNA extraction, cDNA synthesis and qPCR:

Sw.71 cells pellets were lysed and RNA was extracted using the Zymo QuickRNA Microprep kit (Zymo Research, Irvine, CA USA). Whole placentas were equilaterally quartered and homogenized in RNA Lysis buffer and RNA was extracted using the Zymo QuickRNA miniprep kit (Zymo Research, Irvine, CA USA) according to manufacturers recommendations. One microgram (1µg) of placental RNA or Sw.71 cell RNA was converted to cDNA using the High Capacity cDNA synthesis kit (Life Technologies, Carlsbad, CA). cDNA from was diluted 1:20 before use.

Quantitative real-time PCR was carried out using PowerUP SYBR MasterMix (Life Technologies, Carlsbad, CA) and gene-specific primers (Supplementary Table 1) on the ABI 7500 Real-Time PCR System. Cycling conditions were 95℃ for 10 minutes, followed by 40 cycles of 95℃ for 30 secs, 60℃ for 1 minutes. Gene expression was normalized to β-actin and GAPDH, for mouse placenta and human cells, respectively. Fold change was calculated using the delta-delta Ct method. Results are expressed as fold difference from vehicle of same sex. Statistical analysis was performed on dCts.

### Statistical Analysis

For all assays, the dam was considered the experimental unit and tissue from one male and one female per litter was used. For comparisons between Vehicle and CORT-treated animals within a sex (male or female), assays in which the data had a normal distribution (Shapiro-Wilks test) was analyzed using a Student’s T-test. Assays in which the data were not normally distributed were analyzed using a Mann-Whitney Test. Expression analysis in Sw.71 cells treated with Vehicle, CORT and/or RU486 were analyzed using one-way ANOVA. All statistical analysis was conducted using GraphPad Prism.

### Results

In our mouse model of chronic, low-grade maternal stress during pregnancy, offspring born to CORT-treated dams have heightened response to airway introduced allergen, with sex-specific differences in response [9]. Because of the placenta’s well established role as a mediator of sex-specific responses to maternal stress, we evaluated global changes in gene expression in the mouse placenta in response to exogenous stress hormone administration. Pregnant dams were treated with 50µg/mL CORT or vehicle, and whole placenta were collected at E18.5 and RNA-seq analysis was performed (Figure1A). As expected, principal component analysis showed separation by placental sex (Supplementary Figure 1A). Treatment groups were separated on principal component 4, representing 8.7% of the variation in gene expression (Figure 1B). Differential gene expression analysis yielded 238 differentially expressed genes with FDR p-value <0.05 (top 20 DEGs shown in Figure 1C, Supplementary Figure 1B and Supplementary Table 2). 109 genes were upregulated in response to maternal CORT treatment and 129 were downregulated. Interestingly, iodothyronine deiodinase II (*Dio2*) was amongst the most significantly downregulated genes in response to maternal CORT treatment (log_2_(FC) = -1.24, FDR = 2.74×10^-9^, Figure 1D).

Because the highest variance in expression profiles was based on fetal sex, we also analyzed male and female samples separately. In female placenta, 330 DEGs were found (141 upregulated, 189 downregulated; Supplementary Figure 1C and Supplementary Table 3). Interestingly, in male placenta, only 32 genes were differentially expressed in response to maternal CORT treatment (9 up, 23 down; Supplementary Figure 1D and Supplementary Table 4). *Dio2* was significantly downregulated in both sexes however, this downregulation was more robust in female placenta (Figure 1E, log_2_(FC)_female_ = -1.6, p-value_female_ = 3.53 × 10^-15^, FDR_female_ = 5.12 × 10^-11^ ; log_2_(FC)_male_ = -0.8, p-value_male_ = 2.65 × 10^-6^ , FDR_male_ = 0.008).

Next, we validated expression of *Dio2* in whole placenta at E18.5 using qPCR. Consistent with RNA-seq analysis, we observed a downregulation in *Dio2* expression in placenta from CORT-treated dams (Figure 1F-G). There was a greater than two-fold reduction in *Dio2* expression in female placenta from CORT-treated dams, when compared to female controls (p=0.0027, Student’s t-test, Figure 1F). In male placenta from CORT dams, a trend towards decreased *Dio2* expression was observed (Figure 1G).

DIO2, an essential enzyme that converts T4 to T3, the bioactive form of thyroid hormone, is expressed peripheral tissues including the placenta [34]. Thus, we measured TH abundance in E18.5 whole placenta from male and female embryos exposed to Vehicle or CORT treatment *in utero* (Figure 2). There was a 33% reduction in T3 in female placenta from CORT-treated dams when compared to control placenta (p = 0.03, Mann-Whitney Test, Figure 2A). There was only an 11% reduction in T3 in male placenta from CORT-treated dams compared to Vehicle counterparts, and this reduction did not reach statistical significance (Figure 2B). T4 levels were not changed in male or female placenta in response to maternal CORT treatment (Figure 2C-D).

**Figure 2:**
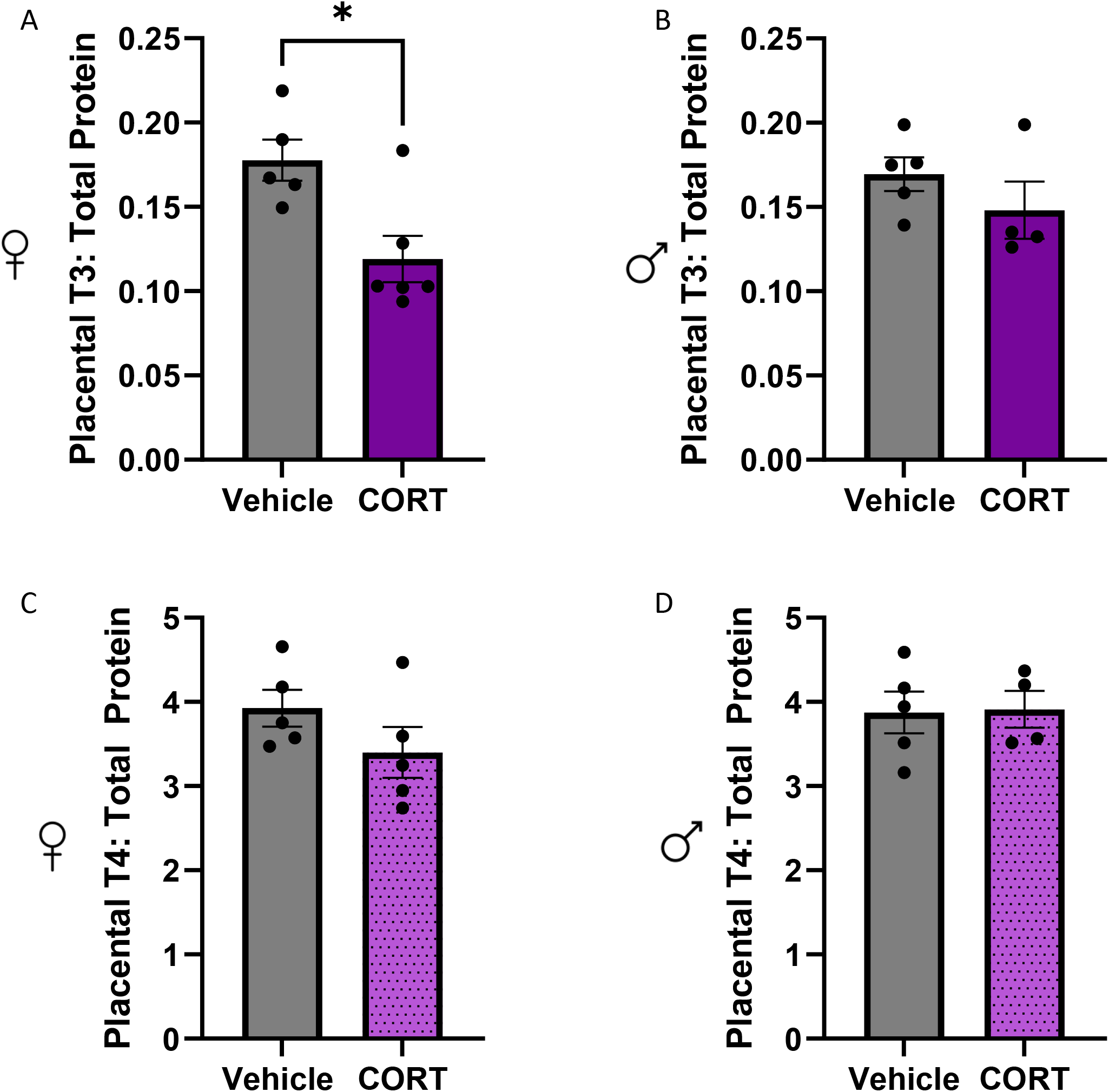
Maternal CORT treatment reduces T3 levels in female placenta at E18.5. A,B) Ratio of T3 (ng/mL) to total placental protein (µg/mL) in females (A) and males (B). C,D) Ratio of T4 (ng/mL) to total placental protein (µg/mL) in females (C) and males (D). n = 4 – 6 placenta/treatment from independent dams. *p = 0.03, Mann-Whitney Test.

During gestation, the placenta mediates the metabolism (T4 to T3) and transport of THs from maternal to fetal circulation [36]. Given the reduction in placental *Dio2* and T3 in female placenta, we measured the T3 bioavailabilty in fetal serum at E18.5 (Figure 3). In female embryos, circulating T3 was significantly reduced in response to prenatal CORT treatment (Figure 3A, p = 0.02, Mann-Whitney Test). In contrast, maternal CORT treatment did not affect circulating T3 levels in male embryos (Figure 3B). To determine whether the female-specific reduction in T3 was due to a reduction in maternal thyroid hormone, TH levels were measured in pregnant dams. T3 and T4 levels were not different between pregnant dams in the CORT-treated group compared to the Vehicle-treated group indicating that the changes in both placental and fetal T3 of female embryos were independent of changes in maternal THs levels at E18.5 (Figure 3C –D). Notably, female embryos from CORT-treated dams were significantly smaller than control counterparts (Supplementary Figure 2A). Placental weights were not affected by maternal CORT treatment or sex (Supplementary Figure 2B).

**Figure 3:**
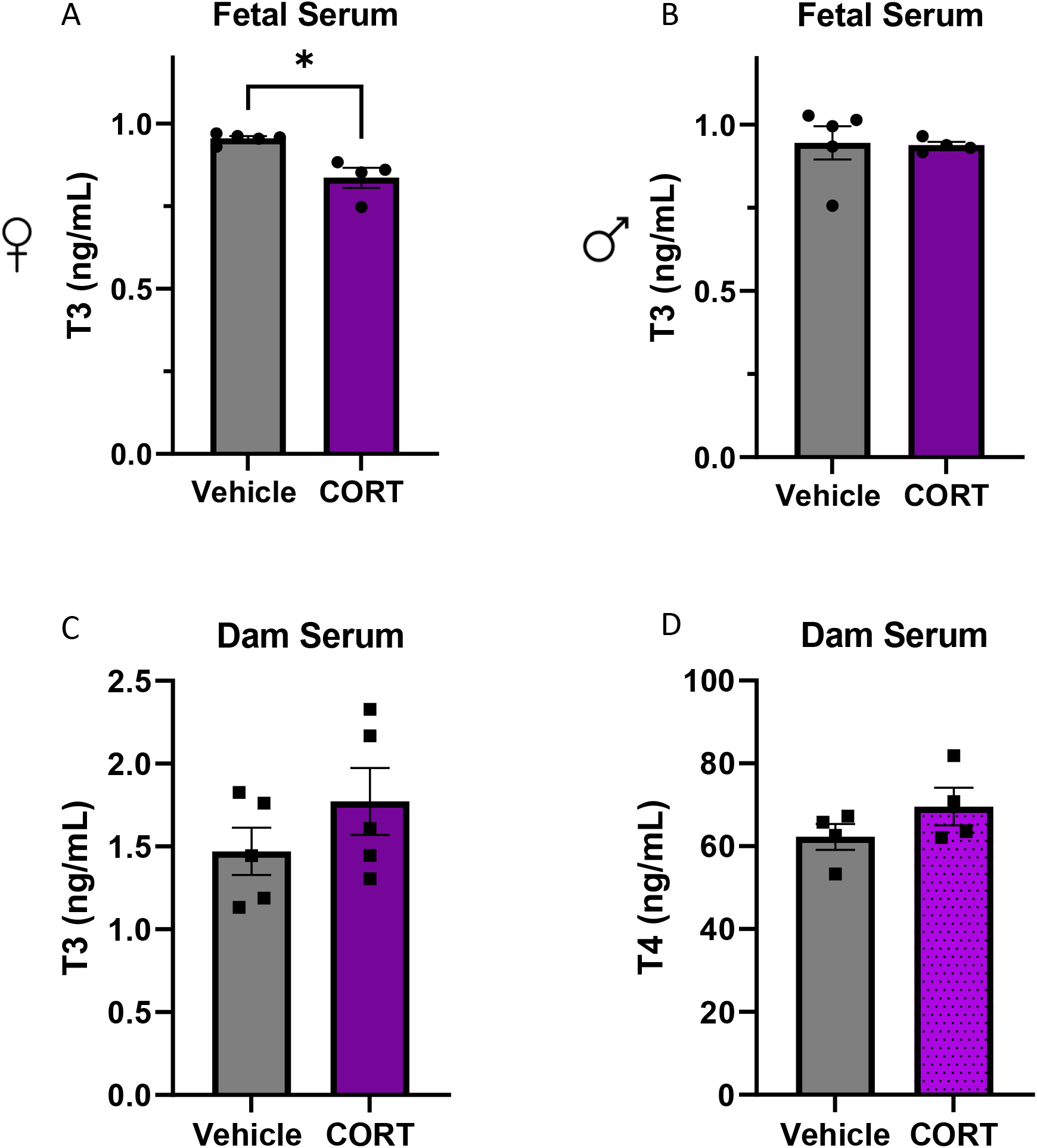
Circulating thyroid hormones in the fetus and dam at E18.5. A) T3 concentration in serum from female embryos at E18.5. B) T3 concentration in serum from male embryos at E18.5. Serum was pooled across embryos of the same sex within a litter. N = 4-5 litters/treatment. *p = 0.02, Mann-Whitney Test. C) T3 concentration in maternal (dam) serum. D) T4 concentration in maternal serum. N = 4 dams/treatment.

In the mouse, the fetal thyroid gland assumes production of thyroid hormones at approximately E17 [41]. Before then, the fetus is dependent on maternally derived THs that are metabolized and/or transported across the placenta. At E16.5, *Dio2* expression was reduced in whole placenta from female and male fetuses in response to maternal, prenatal CORT exposure however, these differences were not statistically significant. (Figure 4 A-B). Additionally, there was no difference in T3 or T4, with respect to sex or maternal treatment, in whole placenta at E16.5 (Figure 4 C-F). Due to low sample yield, we were unable to measure circulating THs in embryos at E16.5. Consistent with these data, we saw no difference in fetal or placental wet weights between male and female from CORT-treated dams when compared to control counterparts (Supplementary Figure 2C - D).

**Figure 4:**
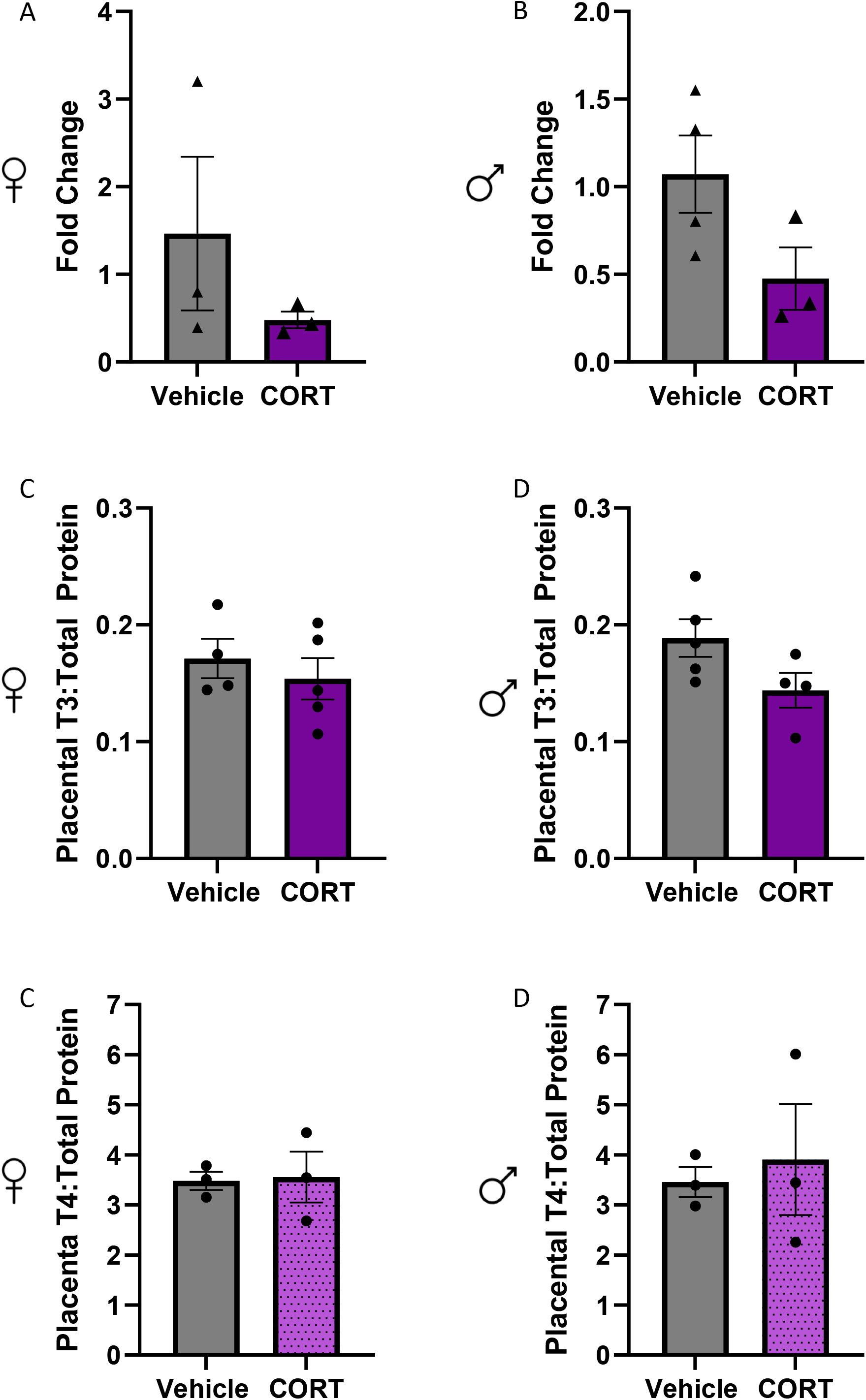
Maternal CORT treatment does not alter thyroid hormone levels in whole placenta at E16.5. A,B) DIO2 expression in female (A) and male (B) whole placenta. N = 3 placenta from independent dams. Gene expression was normalized to beta-actin and set relative to the Vehicle group. Data are presented as average fold change (2^-ddCt) +/-SEM. Statistical analysis was performed on dCt values (see methods). Ratio of T3 (ng/mL) to total placental protein (ug/mL) in females (A) and males (B). C,D) Ratio of T4 (ng/mL) to total placental protein (ug/mL) in females (C) and (D). n = 3 – 5 placenta/treatment from independent dams.

Whole placenta is composed of both fetal (trophoblast) and maternal (decidua) tissues. We hypothesized that maternally derived *Dio2* and THs may mask CORT-induced deficits in placental THs at earlier time points. To evaluate the effects of maternal, prenatal CORT exposure on TH processing and bioavailability in the fetal placenta before the onset of fetal TH production, whole placenta were harvested at E16.5 and the maternal decidua was dissected away. Prenatal CORT treatment resulted in a significant downregulation of *Dio2* expression in trophoblasts from female embryos (Figure 5A, p = 0.01, Unpaired t-test). In male embryos, *Dio2* was not significantly downregulated in trophoblasts in response to maternal CORT treatment (Figure 5B). Consistent with these data, female trophoblasts, but not male, exposed to maternal, prenatal CORT had significantly reduced levels of T3 (Figure 5C-D, p = 0.02, Mann-Whitney Test). Interestingly, we also observed a significant decrease in trophoblast T4 in female tissues from CORT-treated dams, when compared to female controls (Figure 5E, p = 0.03, Mann-Whitney Test). There was no change in T4 in response to maternal CORT in trophoblast from male embryos (Figure 5F). The female-specific changes in trophoblast TH concentrations were independent of changes in circulating THs in pregnant dams at E16.5 (Figure 5G-H).

**Figure 5:**
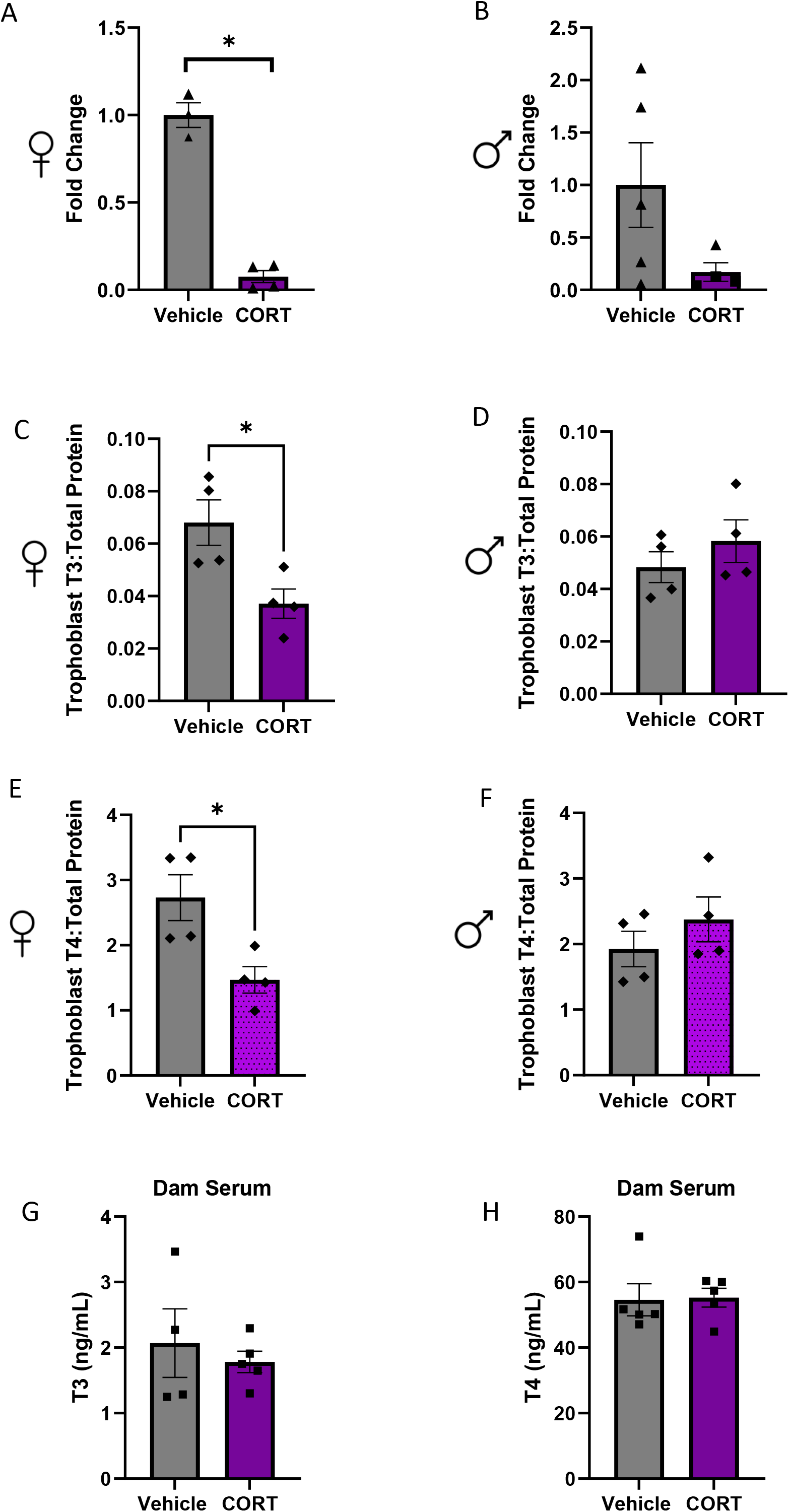
Maternal CORT treatment alters thyroid hormone levels in trophoblasts from female embryos at E16.5. Whole placenta were harvested at E16.5 from Vehicle and CORT-treated dams and the maternal decidua was dissected away. The remaining fetal placental tissue is termed “trophoblast.” A-B) DIO2 expression in female (A) and male (B) trophoblasts. N = 3-5 placenta from independent dams. Gene expression was normalized to beta-actin and set relative to the Vehicle group. Data are presented as average fold change (2^-ddCt) +/-SEM. Statistical analysis was performed on dCt values (see methods). C-D) Ratio of T3 (ng/mL) to total trophoblast protein (ug/mL) in females (C) and males (D). N = 4 trophoblast samples/treatment from independent dams. E-F) Ratio of T4 (ng/mL) to total lacental protein (ug/mL) in females (C) and (D). N = 4 trophoblast samples/treatment from independent dams. *p <0.03, Mann-Whitney Test). G-H) Circulating thyroid hormones in pregnant dams at E16.5. G) T3 concentration in serum from Vehicle and CORT treated dams. H) T4 concentration in serum from Vehicle and CORT treated dams. N = 4 dams/treatment.

To explore the molecular mechanism underlying the CORT-induced downregulation of *Dio2 in vitro*, we utilized Sw.71 cells, a first-trimester extravillous trophoblast cell line [42]. *DIO2* is highly expressed in this cell type and thus makes a good model for exploration [43]. Treatment with 500ng/mL CORT for 6 or 24 hours resulted in a significant decrease in *DIO2* expression (Figure 6A and B, p < 0.01, ANOVA with Tukey), a result consistent with our *in vivo* findings. CORT mediates the stress response by binding to its nuclear receptor, the glucocorticoid receptor (GR), to regulate expression of target genes [18]. To determine if the CORT-induced downregulation of *DIO2* was dependent on the GR, we co-treated cells with a well-documented GR antagonist, RU486. Co-treatment with 500ng/mL CORT and 1uM RU486 for 6hrs or 24hrs was sufficient to rescue expression of DIO2 to control levels in Sw.71 cells (Figure 6).

**Figure 6:**
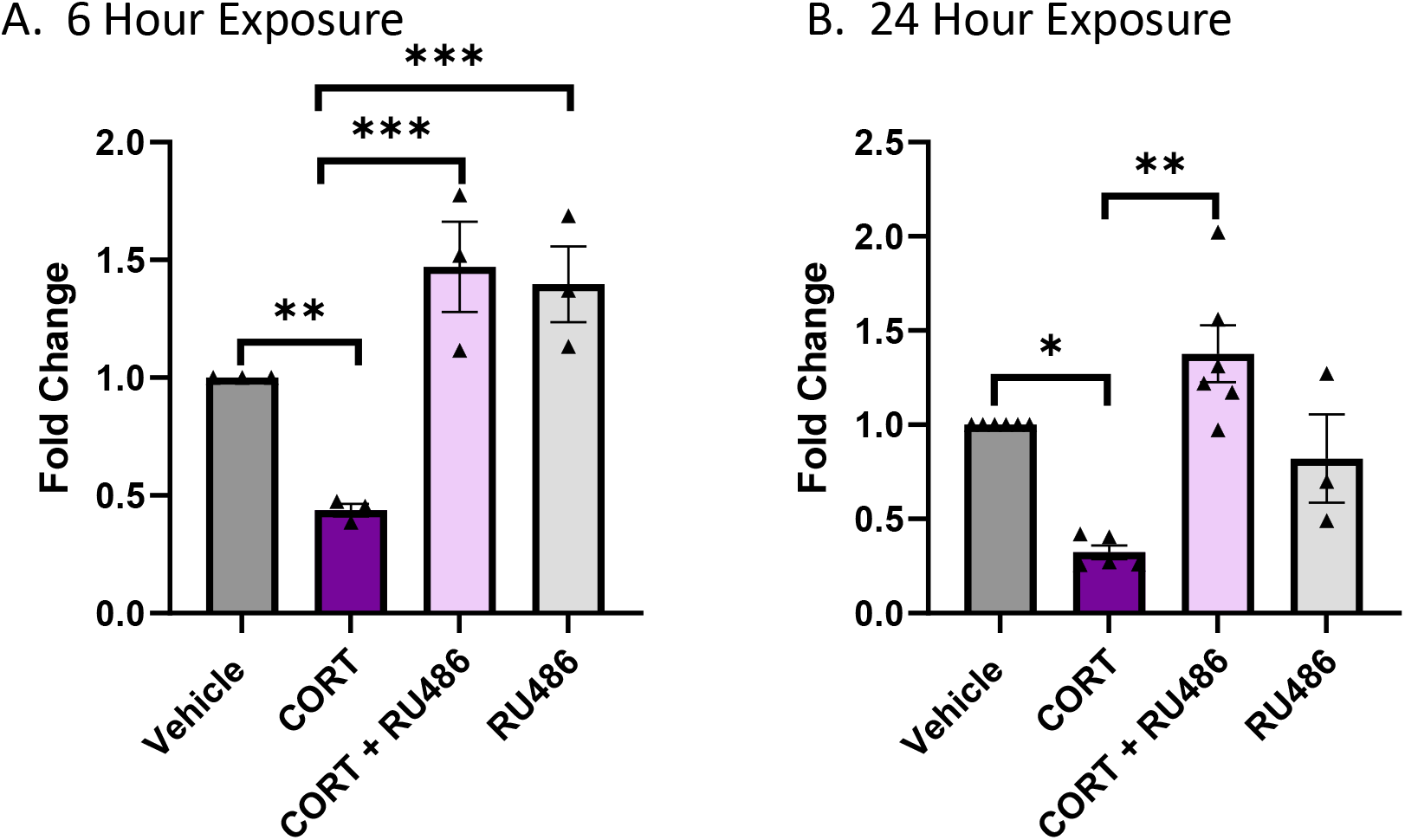
CORT treatment downregulates *DIO2* in an extravillous trophoblast cell line. Sw.71 cells were treated with Vehicle, 500ng/mL CORT, 500ng/mL CORT with 1uM RU486 or 1uM RU486 for A) 6 hours (N = 3 independent replicates) or B) 24 hours (N = 3-6 independent replicates). DIO2 expression was normalized to GAPDH and set relative to the Vehicle group. Data are presented as average fold change (2^-ddCt) +/-SEM. Statistical analysis was performed on dCt values (see methods), *p=0.01, **p<0.005, ***p<0.0001

## Discussion

In this study, we used a recently reported mouse model to identify cellular and molecular pathways in the placenta that are sensitive to maternal, prenatal stress. In this model, pregnant dams receiving exogenous stress hormone have an approximate two-fold increase in circulating CORT levels, without a concomitant increase in fetal circulating CORT at the end of gestation [9]. As the placenta is an intermediary between the maternal environment and the fetus, this mouse model is well suited for examination of placenta-mediated responses to excess maternal stress hormone. Here we show that exogenous administration of low-dose CORT to mice during the second half of pregnancy reduces expression of a key TH biosynthesis gene, *Dio2*, in the placenta and alters TH availability in the placenta and fetus in a sex-specific manner. To the best of our knowledge, this is the first work to consider the effects of maternal CORT excess on placental TH production and bioavailability during gestation in the mouse.

### Maternal CORT alters *DIO2* expression in placenta

Our data suggest that CORT regulates expression of *DIO2*. In the mouse placenta, maternal treatment with exogenous CORT during pregnancy reduced *Dio2* expression at both E16.5, before the onset of fetal TH production, and at E18.5, after fetal TH production commences [41]. CORT treatment also reduced expression of *DIO2* in a human extravillous trophoblast cell line. These results suggest evolutionary conservation of CORT-mediated downregulation of *DIO2* in the placenta. Consistent with this, glucocorticoid-induced downregulation of *Dio2* has also been observed in other tissues [44]. Interestingly, in aortic media of rats, *Dio2* expression and activity was inversely related to the diurnal CORT surge, with expression being lowest when plasma CORT levels were high [45].

The CORT-induced downregulation of *DIO2* in the placenta may serve a physiological role. In the mouse, as gestation progresses, maternal CORT synthesis and passage across the placenta increases, largely to support maturation of fetal organ systems. Concomitantly, the fetus begins to synthesize their own thyroid hormone [36]. It is possible that the rise in CORT downregulates *Dio2* as the burden for THs during gestations switches from the dam and placenta to the fetus. While similar phenomenon exists in the human placenta, there are species-specific differences in the timing of fetal thyroid hormone production. The human fetus produces its own thyroid hormone as early as the second trimester (∼20 weeks) whereas this transition happens near term in the mouse [34]. More work must be done to unravel the spatiotemporal expression and activity patterns of Dio2 in the mouse placenta, and in response to CORT, across gestation.

We used Sw.71 cells, a human extravillous trophoblast cell line, to explore the molecular mechanisms underlying the CORT-induced downregulation of *DIO2*. Single cell sequencing analysis has identified *DIO2* as a marker of the extravillous trophoblast lineage in human placenta in early pregnancy [46], making it a good model for this exploration. We found that downregulation of *DIO2* by CORT was dependent on the GR. CORT-bound GR complexes regulate gene expression by binding glucocorticoid response elements (GREs) in gene regulatory regions [18]. Functional GREs in the *DIO2* promoter region have been reported [47]. Further, we identified three putative GRE binding sites upstream of the human DIO2 promoter (UCSC Genome Browser, data not shown). These data raise the possibility that the CORT-mediated downregulation of *DIO2* in placenta is the result of direct inhibition of gene expression at the genomic level.

### Sex-specific differences in placental/fetal TH following maternal CORT treatment

T3 concentrations were reduced in the placenta and in the fetus in response to maternal CORT treatment, but only in tissues and serum from female fetuses. Maternal THs are metabolized and transported across the mouse placenta throughout gestation [36]. Two DIO – family proteins catalyze the conversion of T4 into T3, Dio1 and Dio2. Dio1 is responsible for circulating T3 levels and is highly expressed in the liver and kidneys. Dio1 is lowly expressed in the placenta, and likely makes only modest contributions to placentally derived T3. DIO2, in contrast, regulates T3 production in peripheral tissue and is expressed in the placenta [48]. In vascular smooth muscle cells and adipose tissue of rats, glucocorticoids reduced both expression of *DIO2* at the mRNA level and DIO2 activity [45, 47]. Additionally, cortisol treatment of human term placental cells reduced the rate of T4 deiodination, the process by which T3 is generated [49]. Thus, the observed reduction in T3 in placenta and fetuses from CORT-treated dams is consistent with the observed downregulation in *DIO2*.

The reduction of T3 in females may be driven by CORT mediated, sex-specific changes in placental gene expression. The placenta is recognized as a sexually dimorphic organ, with sex chromosome genes being differentially expressed and driving basal differences in placental function and response based on sex [6]. In fact, prenatal glucocorticoid excess can lead to sex-specific changes in expression of the GR itself in the placenta [20, 50]. In pregnant mice exposed to acute sound stress, female fetuses displayed increased serum CORT, but not males. In this model, *11β-HSD2* and ABC transporters that function to regulate CORT efflux into maternal circulation, were elevated in male placenta but not female. Consistent with this, female offspring displayed more severe fetal growth restriction than male offspring born to stressed dams [51]. In the work presented here, reductions in *Dio2* expression in placental tissues and T3 levels in serum, were more robust in females than in males. It is possible that this is due to sex-specific differential regulation of the *Dio2* gene by the GR. It is also possible that other key TH biosynthesis and transport genes are altered in a sex-specific manner and accounts for the female-specific molecular phenotype observed here.

### Teasing apart maternal vs. fetal placental contribution to thyroid hormone misregulation by CORT

At E18.5, we observed decreased expression of *Dio2* and female-specific reduction of T3 in whole placenta. In contrast, at E16.5, while there was a reduction in *Dio2* expression, there was no difference in T3 levels in whole placenta in response to maternal CORT treatment. However, when the maternal decidua was removed from E16.5 placenta, both *Dio2* expression and T3 levels were reduced in the fetal portion (trophoblasts) of female placenta. Notably, T4 was also reduced in female trophoblasts, a result not observed in whole placenta at E18.5. Zone-specific responses to maternal prenatal insults have been reported [22, 52]. In rats fed a high fat diet, reductions in genes involved TH biosynthesis and signaling pathway, including *Dio2*, were restricted to the fetal-derived junctional zone at E18.5 [52]. Up until E17, the mouse fetus is dependent on maternally derived thyroid hormones and thyroid hormone in the maternal compartment far exceeds that found in the fetal spaces [41]. These data suggest that, at E16.5, the maternal decidua is a sink for thyroid hormones, which may be transported across the placenta to supply the fetus. This interpretation is complicated, however, by the lack of zone-specific (maternal vs. fetal) analysis at E18.5.

### Consequences of reduced fetal T3

With the mouse model of chronic maternal stress used here, we previously reported that female offspring born to CORT-treated dams have increased mucus cell metaplasia in response to airway introduced allergen (HDM), when compared to female control counterparts [9]. It is possible that this phenotype is driven by CORT-induced impairment of lung epithelial morphogenesis in female mice, who already display marked differences in fetal lung development and sensitivity to maternal prenatal insults [11, 53]. Notably, TH signaling during pregnancy is important for development and maturation of the fetal lung [54, 55]. It is possible that the female-specific reduction in T3 alters fetal lung development and predisposes to enhance mucus cell metaplasia. Further work must be done to confirm such a mechanism.

Approximately 80% of the TH secreted by the thyroid gland is T4. *DIO2* catalyzes the conversion of this T4 to T3 in tissues, including the placenta [36]. Interestingly, a recent report suggests that in human trophoblasts cells, *DIO2* functions to regulate trophoblast proliferation, invasion and migration, in a manner independent of TH metabolism [37]. Thus, the consequence of CORT-induced downregulation of placental *DIO2* on trophoblast/placental function should be explored further.

### Maternal stress and disruption of fetal thyroid axis

In pregnant ewes, prenatal dexamethasone treatment increased T3, but not T4 concentrations, in the fetus and decreased TH in maternal circulation [29]. While these data offer precedence for a relationship between misregulated maternal glucocorticoids and the fetal thyroid axis, they are in contrast to the data presented here where maternal TH concentration was unaffected by exogenous CORT during pregnancy and fetal T3 was decreased specifically in female fetuses. However, these differences in results may reflect differences in the model organism used and the timing and type of glucocorticoid excess. Specifically, dexamethasone is poorly metabolized by 11β-HSD2 and readily crosses the placenta to affect fetal growth in a direct manner [56]. Our model is unique in that very little of the excess maternal CORT reaches the fetus and thus, we hypothesize that the observed fetal effects are largely mediated by the placenta.

In summary, elevated maternal stress hormone during late pregnancy alters gene expression in the placenta. Specifically, prenatal exposure to CORT alters thyroid hormone synthesis and bioavailability in the mouse placenta and fetus, with female offspring being particularly affected. This work adds to a body of evidence supporting a role for maternal stress, and other mechanisms of glucocorticoid excess (i.e clinical administration of dexamethasone in threatened pre-term birth) on misregulation of THs in pregnancy [29, 31].

## Supporting information

Suppmentary Table

## Abbreviations

CORT: corticosterone (mouse) or cortisol (human)
TH: thyroid hormones (T3 and T4)
T3: triiodothyronine
T4: thyroxine
E: embryonic day

## Author contributions

ENP, SS and RR performed and analyzed experiments, and aided in manuscript preparation. IR and LV performed and analyzed experiments. SA aided in data analysis. KER developed project idea, oversaw design and performance of experiments, performed data and statistical analysis and aided in the preparation of the manuscript. ALS developed project ideas, oversaw design and performance of experiments, performed data and statistical analysis and wrote the manuscript.

## Acknowledgements

We would like to acknowledge the Van Andel Research Institutes’ Genomic Core. We also gratefully acknowledge the animal facility staff at the Van Andel Research Institute Vivarium, at the Michigan State University College of Human Medicine and at Kenyon College.

## Supplementary Figure Legends

**Supplementary Figure 1: Differential gene expression in placenta from Vehicle and CORT-treated dams at E18.5.** A) Principal component analysis of placental transcriptomes by sex (pink dots represent female samples, blue dots represent male Samples). B) Heatmap of top 100 differentially expressed genes at FDR <0.05. n = 8 placenta from 4 dams/treatment – one male and one female placenta were selected from a single dam. C) Volcano plot showing 330 differentially expressed genes (red) at a false discovery rate adjusted p-value <0.05 in females. Dio2 is indicated. D) Volcano plot showing 32 differentially expressed genes (red) at a false discovery rate adjusted p-value <0.05 in males. Dio2 is indicated.

**Supplementary Figure 2: Maternal CORT treatment reduces fetal weight of female embryos at E18.5.** Fetal wet weights at E18.5 (A) and E16.5 (E) were taken at the time of collection. Placenta wet weights at E18.5 (B) and E16.5 (F) were taken at the time of collection. Placental efficiency at E18.5 (C) and E16.5 (D) were calculated by dividing fetal weight (in grams) by placental weight (in grams). Number of embryos per litter at E18.5 (D) and E16.5 (H). N at E18.5 = 15 – 22 embryos from 4-5 litters. N at E16.5 = 11-17 embryos from 7-8 litters. *p = 0.01, **p = 0.009.

## References

[1] A.L. Fowden, Stress during pregnancy and its life-long consequences for the infant, J Physiol 595(15) (2017) 5055–5056.

[2] E.J. Mulder, P.G. Robles de Medina, A.C. Huizink, B.R. Van den Bergh, J.K. Buitelaar, G.H. Visser, Prenatal maternal stress: effects on pregnancy and the (unborn) child, Early Hum Dev 70(1-2) (2002) 3–14.

[3] P.H. Rondo, R.F. Ferreira, F. Nogueira, M.C. Ribeiro, H. Lobert, R. Artes, Maternal psychological stress and distress as predictors of low birth weight, prematurity and intrauterine growth retardation, Eur J Clin Nutr 57(2) (2003) 266–72.

[4] E.C. Cottrell, J.R. Seckl, Prenatal stress, glucocorticoids and the programming of adult disease, Front Behav Neurosci 3 (2009) 19.

[5] T.L. Bale, Sex differences in prenatal epigenetic programming of stress pathways, Stress 14(4) (2011) 348–56.

[6] T.L. Bale, The placenta and neurodevelopment: sex differences in prenatal vulnerability, Dialogues Clin Neurosci 18(4) (2016) 459–464.

[7] K.M. Schulz, J.N. Pearson, E.W. Neeley, R. Berger, S. Leonard, C.E. Adams, K.E. Stevens, Maternal stress during pregnancy causes sex-specific alterations in offspring memory performance, social interactions, indices of anxiety, and body mass, Physiol Behav 104(2) (2011) 340–7.

[8] A.L. Smith, P. Bole Aldo, K.E. Racicot, Chapter 17 - Placental regulation of immune functions, in: G. Mor (Ed.), Reproductive Immunology, Academic Press2021, pp. 335–348.

[9] A.L. Smith, E. Paul, D. McGee, R. Sinniah, E. Flom, D. Jackson-Humbles, J. Harkema, K.E. Racicot, Chronic, Elevated Maternal Corticosterone During Pregnancy in the Mouse Increases Allergic Airway Inflammation in Offspring, Front Immunol 10 (2019) 3134.

[10] S. Sutherland, S.M. Brunwasser, Sex Differences in Vulnerability to Prenatal Stress: a Review of the Recent Literature, Curr Psychiatry Rep 20(11) (2018) 102.

[11] D.E. Zazara, M. Wegmann, A.D. Giannou, A.M. Hierweger, M. Alawi, K. Thiele, S. Huber, M. Pincus, A.C. Muntau, M.E. Solano, P.C. Arck, A prenatally disrupted airway epithelium orchestrates the fetal origin of asthma in mice, J Allergy Clin Immunol 145(6) (2020) 1641–1654.

[12] G.J. Burton, A.L. Fowden, The placenta: a multifaceted, transient organ, Philos Trans R Soc Lond B Biol Sci 370(1663) (2015) 20140066.

[13] K. Chapman, M. Holmes, J. Seckl, 11beta-hydroxysteroid dehydrogenases: intracellular gate-keepers of tissue glucocorticoid action, Physiol Rev 93(3) (2013) 1139–206.

[14] L.I. Stirrat, B.G. Sengers, J.E. Norman, N.Z.M. Homer, R. Andrew, R.M. Lewis, R.M. Reynolds, Transfer and Metabolism of Cortisol by the Isolated Perfused Human Placenta, J Clin Endocrinol Metab 103(2) (2018) 640–648.

[15] H.J. Speirs, J.R. Seckl, R.W. Brown, Ontogeny of glucocorticoid receptor and 11beta-hydroxysteroid dehydrogenase type-1 gene expression identifies potential critical periods of glucocorticoid susceptibility during development, J Endocrinol 181(1) (2004) 105–16.

[16] L.A. Welberg, K.V. Thrivikraman, P.M. Plotsky, Chronic maternal stress inhibits the capacity to up-regulate placental 11beta-hydroxysteroid dehydrogenase type 2 activity, J Endocrinol 186(3) (2005) R7–R12.

[17] Z. Saif, N.A. Hodyl, E. Hobbs, A.R. Tuck, M.S. Butler, A. Osei-Kumah, V.L. Clifton, The human placenta expresses multiple glucocorticoid receptor isoforms that are altered by fetal sex, growth restriction and maternal asthma, Placenta 35(4) (2014) 260–8.

[18] S. Timmermans, J. Souffriau, C. Libert, A General Introduction to Glucocorticoid Biology, Front Immunol 10 (2019) 1545.

[19] V.L. Clifton, J. Cuffe, K.M. Moritz, T.J. Cole, P.J. Fuller, N.Z. Lu, S. Kumar, S. Chong, Z. Saif, Review: The role of multiple placental glucocorticoid receptor isoforms in adapting to the maternal environment and regulating fetal growth, Placenta 54 (2017) 24–29.

[20] J.S. Cuffe, L. O’Sullivan, D.G. Simmons, S.T. Anderson, K.M. Moritz, Maternal corticosterone exposure in the mouse has sex-specific effects on placental growth and mRNA expression, Endocrinology 153(11) (2012) 5500–11.

[21] B. Baisden, S. Sonne, R.M. Joshi, V. Ganapathy, P.S. Shekhawat, Antenatal dexamethasone treatment leads to changes in gene expression in a murine late placenta, Placenta 28(10) (2007) 1082–90.

[22] J.Y. Lee, H.J. Yun, C.Y. Kim, Y.W. Cho, Y. Lee, M.H. Kim, Prenatal exposure to dexamethasone in the mouse induces sex-specific differences in placental gene expression, Dev Growth Differ 59(6) (2017) 515–525.

[23] B.R. Mueller, T.L. Bale, Sex-specific programming of offspring emotionality after stress early in pregnancy, J Neurosci 28(36) (2008) 9055–65.

[24] H. Shang, L. Sun, T. Braun, Q. Si, J. Tong, Revealing the action mechanisms of dexamethasone on the birth weight of infant using RNA-sequencing data of trophoblast cells, Medicine (Baltimore) 97(4) (2018) e9653.

[25] C.L. Howerton, T.L. Bale, Targeted placental deletion of OGT recapitulates the prenatal stress phenotype including hypothalamic mitochondrial dysfunction, Proc Natl Acad Sci U S A 111(26) (2014) 9639–44.

[26] C.L. Howerton, C.P. Morgan, D.B. Fischer, T.L. Bale, O-GlcNAc transferase (OGT) as a placental biomarker of maternal stress and reprogramming of CNS gene transcription in development, Proc Natl Acad Sci U S A 110(13) (2013) 5169–74.

[27] S.L. Bronson, T.L. Bale, The Placenta as a Mediator of Stress Effects on Neurodevelopmental Reprogramming, Neuropsychopharmacology 41(1) (2016) 207–18.

[28] E.P. Davis, C.A. Sandman, The timing of prenatal exposure to maternal cortisol and psychosocial stress is associated with human infant cognitive development, Child Dev 81(1) (2010) 131–48.

[29] A.J. Forhead, J.K. Jellyman, D.S. Gardner, D.A. Giussani, E. Kaptein, T.J. Visser, A.L. Fowden, Differential effects of maternal dexamethasone treatment on circulating thyroid hormone concentrations and tissue deiodinase activity in the pregnant ewe and fetus, Endocrinology 148(2) (2007) 800–5.

[30] M.T. Ruis, K.D. Rock, S.M. Hall, B. Horman, H.B. Patisaul, H.M. Stapleton, PBDEs Concentrate in the Fetal Portion of the Placenta: Implications for Thyroid Hormone Dysregulation, Endocrinology 160(11) (2019) 2748–2758.

[31] S. St-Cyr, S. Abuaish, K.C. Welch, Jr., P.O. McGowan, Maternal predator odour exposure programs metabolic responses in adult offspring, Sci Rep 8(1) (2018) 8077.

[32] M.A. Suter, H. Sangi-Haghpeykar, L. Showalter, C. Shope, M. Hu, K. Brown, S. Williams, R.A. Harris, K.L. Grove, R.H. Lane, K.M. Aagaard, Maternal high-fat diet modulates the fetal thyroid axis and thyroid gene expression in a nonhuman primate model, Mol Endocrinol 26(12) (2012) 2071–80.

[33] C. Luongo, M. Dentice, D. Salvatore, Deiodinases and their intricate role in thyroid hormone homeostasis, Nat Rev Endocrinol 15(8) (2019) 479–488.

[34] A.J. Forhead, A.L. Fowden, Thyroid hormones in fetal growth and prepartum maturation, J Endocrinol 221(3) (2014) R87–R103.

[35] M.D. Kilby, J. Verhaeg, N. Gittoes, D.A. Somerset, P.M. Clark, J.A. Franklyn, Circulating thyroid hormone concentrations and placental thyroid hormone receptor expression in normal human pregnancy and pregnancy complicated by intrauterine growth restriction (IUGR), J Clin Endocrinol Metab 83(8) (1998) 2964–71.

[36] A. Eerdekens, J. Verhaeghe, V. Darras, G. Naulaers, G. Van den Berghe, L. Langouche, C. Vanhole, The placenta in fetal thyroid hormone delivery: from normal physiology to adaptive mechanisms in complicated pregnancies, J Matern Fetal Neonatal Med 33(22) (2020) 3857–3866.

[37] E.A. Adu-Gyamfi, J. Lamptey, X.M. Chen, F.F. Li, C. Li, L.L. Ruan, X.N. Yang, T.H. Liu, Y.X. Wang, Y.B. Ding, Iodothyronine deiodinase 2 (DiO2) regulates trophoblast cell line cycle, invasion and apoptosis; and its downregulation is associated with early recurrent miscarriage, Placenta 111 (2021) 54–68.

[38] E. Vasilopoulou, L.S. Loubiere, H. Heuer, M. Trajkovic-Arsic, V.M. Darras, T.J. Visser, G.E. Lash, G.S. Whitley, C.J. McCabe, J.A. Franklyn, M.D. Kilby, S.Y. Chan, Monocarboxylate transporter 8 modulates the viability and invasive capacity of human placental cells and fetoplacental growth in mice, PLoS One 8(6) (2013) e65402.

[39] D. Qu, A. McDonald, K.J. Whiteley, S.A. Bainbridge, S.L. Adamson, 44 - Layer-Enriched Tissue Dissection of the Mouse Placenta in Late Gestation, in: B.A. Croy, A.T. Yamada, F. J. DeMayo, S.L. Adamson (Eds.), The Guide to Investigation of Mouse Pregnancy, Academic Press, Boston, 2014, pp. 529-535.

[40] J.F. Lambert, B.O. Benoit, G.A. Colvin, J. Carlson, Y. Delville, P.J. Quesenberry, Quick sex determination of mouse fetuses, J Neurosci Methods 95(2) (2000) 127–32.

[41] S. Barez-Lopez, A. Guadano-Ferraz, Thyroid Hormone Availability and Action during Brain Development in Rodents, Front Cell Neurosci 11 (2017) 240.

[42] S.L. Straszewski-Chavez, V.M. Abrahams, A.B. Alvero, P.B. Aldo, Y. Ma, S. Guller, R. Romero, G. Mor, The isolation and characterization of a novel telomerase immortalized first trimester trophoblast cell line, Swan 71, Placenta 30(11) (2009) 939–48.

[43] M. Karlsson, C. Zhang, L. Mear, W. Zhong, A. Digre, B. Katona, E. Sjostedt, L. Butler, J. Odeberg, P. Dusart, F. Edfors, P. Oksvold, K. von Feilitzen, M. Zwahlen, M. Arif, O. Altay, X. Li, M. Ozcan, A. Mardinoglu, L. Fagerberg, J. Mulder, Y. Luo, F. Ponten, M. Uhlen, C. Lindskog, A single -cell type transcriptomics map of human tissues, Sci Adv 7(31) (2021).

[44] C.A. Martinez, I. Marteinsdottir, A. Josefsson, G. Sydsjo, E. Theodorsson, H. Rodriguez-Martinez, Prenatal stress, anxiety and depression alter transcripts, proteins and pathways associated with immune responses at the maternal-fetal interfacedagger, Biol Reprod 106(3) (2022) 449–462.

[45] N. Toyoda, S. Yasuzawa-Amano, E. Nomura, A. Yamauchi, K. Nishimura, C. Ukita, S. Morimoto, A. Kosaki, T. Iwasaka, J.W. Harney, P.R. Larsen, M. Nishikawa, Thyroid hormone activation in vascular smooth muscle cells is negatively regulated by glucocorticoid, Thyroid 19(7) (2009) 755–63.

[46] H. Suryawanshi, P. Morozov, A. Straus, N. Sahasrabudhe, K.E.A. Max, A. Garzia, M. Kustagi, T. Tuschl, Z. Williams, A single-cell survey of the human first-trimester placenta and decidua, Sci Adv 4(10) (2018) eaau4788.

[47] R. Martinez-deMena, R.M. Calvo, L. Garcia, M.J. Obregon, Effect of glucocorticoids on the activity, expression and proximal promoter of type II deiodinase in rat brown adipocytes, Mol Cell Endocrinol 428 (2016) 58–67.

[48] S. Chan, S. Kachilele, E. Hobbs, J.N. Bulmer, K. Boelaert, C.J. McCabe, P.M. Driver, A.R. Bradwell, M. Kester, T.J. Visser, J.A. Franklyn, M.D. Kilby, Placental iodothyronine deiodinase expression in normal and growth-restricted human pregnancies, J Clin Endocrinol Metab 88(9) (2003) 4488–95.

[49] J.T. Hidal, M.M. Kaplan, Inhibition of thyroxine 5’-deiodination type II in cultured human placental cells by cortisol, insulin, 3’, 5’-cyclic adenosine monophosphate, and butyrate, Metabolism 37(7) (1988) 664–8.

[50] J.S.M. Cuffe, Z. Saif, A.V. Perkins, K.M. Moritz, V.L. Clifton, Dexamethasone and sex regulate placental glucocorticoid receptor isoforms in mice, J Endocrinol 234(2) (2017) 89–100.

[51] A. Wieczorek, C.V. Perani, M. Nixon, M. Constancia, I. Sandovici, D.E. Zazara, G. Leone, M.Z. Zhang, P.C. Arck, M.E. Solano, Sex-specific regulation of stress-induced fetal glucocorticoid surge by the mouse placenta, Am J Physiol Endocrinol Metab 317(1) (2019) E109–E120.

[52] J. Saben, P. Kang, Y. Zhong, K.M. Thakali, H. Gomez-Acevedo, S.J. Borengasser, A. Andres, T.M. Badger, K. Shankar, RNA-seq analysis of the rat placentation site reveals maternal obesity-associated changes in placental and offspring thyroid hormone signaling, Placenta 35(12) (2014) 1013–20.

[53] D.R. Curran, L. Cohn, Advances in mucous cell metaplasia: a plug for mucus as a therapeutic focus in chronic airway disease, Am J Respir Cell Mol Biol 42(3) (2010) 268–75.

[54] K. Archavachotikul, T.J. Ciccone, M.R. Chinoy, H.C. Nielsen, M.V. Volpe, Thyroid hormone affects embryonic mouse lung branching morphogenesis and cellular differentiation, Am J Physiol Lung Cell Mol Physiol 282(3) (2002) L359–69.

[55] R. Hume, K. Richard, E. Kaptein, E.L. Stanley, T.J. Visser, M.W. Coughtrie, Thyroid hormone metabolism and the developing human lung, Biol Neonate 80 Suppl 1 (2001) 18–21.

[56] L.S. Kerzner, B.S. Stonestreet, K.Y. Wu, G. Sadowska, M.P. Malee, Antenatal dexamethasone: effect on ovine placental 11beta-hydroxysteroid dehydrogenase type 2 expression and fetal growth, Pediatr Res 52(5) (2002) 706–12.

