## Supplementary figures and images for "Exogenous corticosterone administration during pregnancy alters placental and fetal thyroid hormone availability in females"

### Supplementary Figures

#### Slide 1
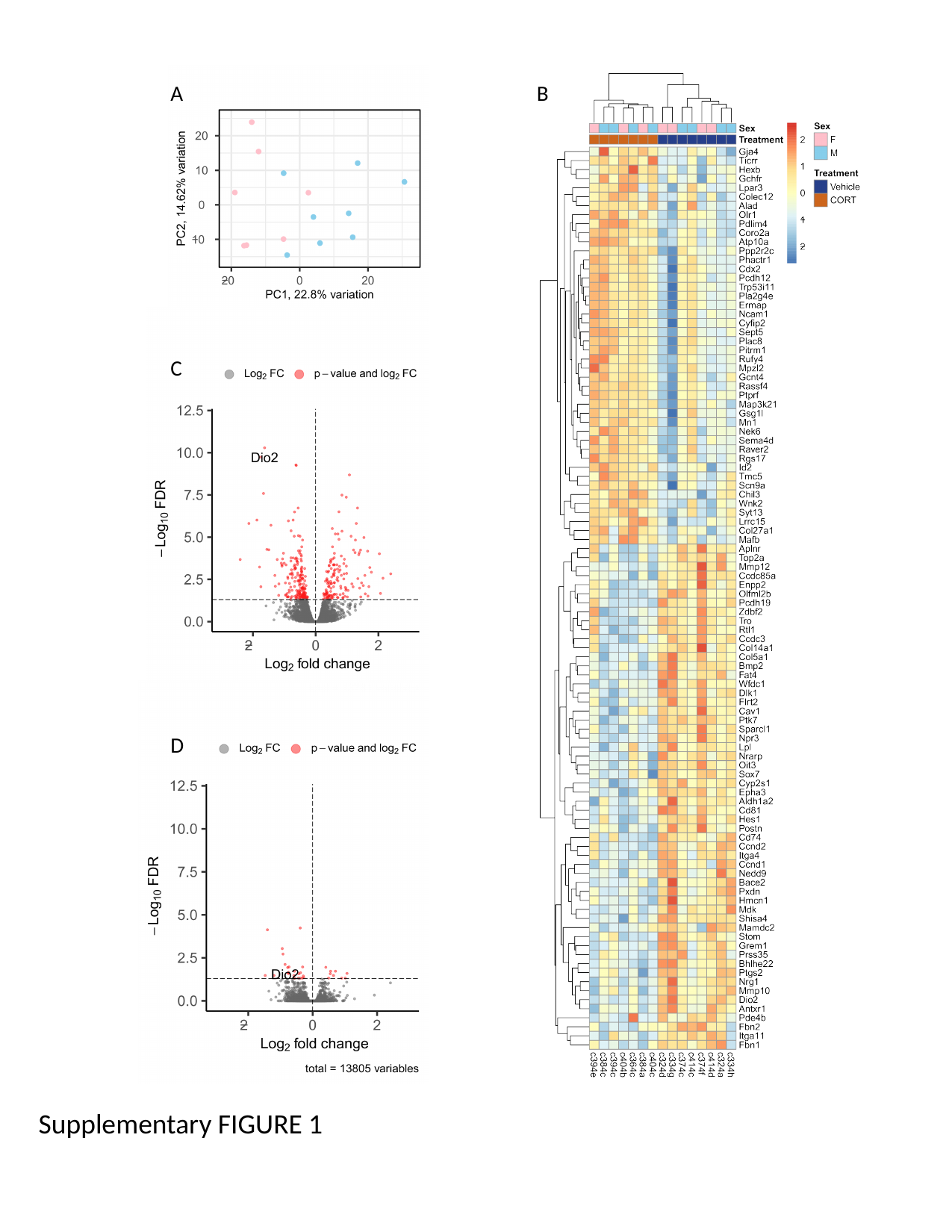

A
B
C
D
Supplementary FIGURE 1

#### Slide 2
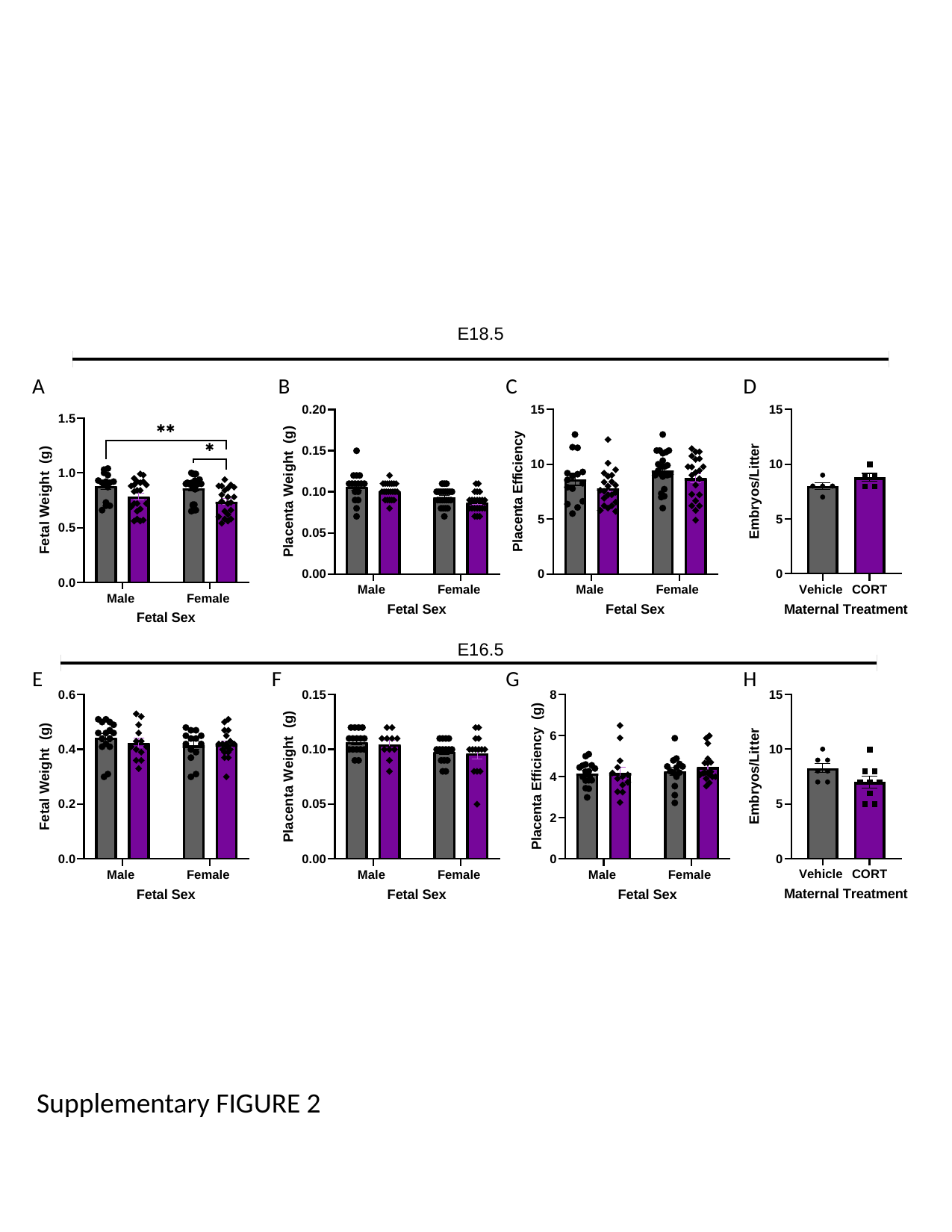

C
D
B
A
G
H
F
E
Supplementary FIGURE 2
